# Inhibition of cell proliferation does not slow down echinoderm neural regeneration

**DOI:** 10.1101/066860

**Authors:** Vladimir S. Mashanov, Olga R. Zueva, José E. García-Arrarás

**Affiliations:** University of Puerto Rico, Rio Piedras, Puerto Rico; University of North Florida, Jacksonville, FL

## Abstract

**Background:** Regeneration of the damaged central nervous system is one of the most interesting post-embryonic developmental phenomena. Two distinct cellular events have been implicated in supplying regenerative neurogenesis with cellular material – generation of new cells through cell proliferation and recruitment of already existing cells through cell migration. The relative contribution and importance of these two mechanisms is often unknown.

**Methods:** Here, we use the regenerating radial nerve cord (RNC) of the echinoderm *Holothuria glaberrima* as a model of extensive post-traumatic neurogenesis in the deuterostome central nervous system. To uncouple the effects of cell proliferation from those of cell migration, we treated regenerating animals with aphidicolin, a specific inhibitor of S-phase DNA replication. Cell migration was tracked with vital staining with the lipophilic dye DiI.

**Results:** Aphidicolin treatment resulted in a significant 2.1-fold decrease in cell proliferation. In spite of this, the regenerating RNC in the treated animals did not differ in histological architecture, size and cell number from its counterpart in the control vehicle-treated animals. DiI labeling showed extensive cell migration in the RNC. Some cells migrated from as far as 2 mm away from the injury plane to contribute to the neural outgrowth.

**Conclusions:** We suggest that inhibition of cell division in the regenerating RNC of H. *glaberrima*, is compensated for by recruitment of cells, which migrate into the RNC outgrowth from deeper regions of the neuroepithelium. Neural regeneration in echinoderms is thus a highly regulative developmental phenomenon, in which the size of the cell pool can be controlled either by cell proliferation or cell migration, and the latter can neutralize perturbations in the former.

## 1 Introduction

Echinoderms are well known for their outstanding capacity to regenerate parts of their body, including the central nervous system (CNS), in response to injury or autotomy (Candia Carnevali, 2006). The cellular and molecular events that allow echinoderms to restore the integrity of their organs and appendages with such efficiency have been a major focus in echinoderm regenerative biology. It has been established that the earliest response to injury involves extensive dedifferentiation of adult tissues, when mature cells greatly simplify their shape and internal organization through elimination of specialized cytoplasmic structures (Candia Carnevali, 2006; García-Arrarás and Dolmatov, 2010; Mashanov et al., 2013). Dedifferention primes the adult tissues for the subsequent morphogenetic phase, which in turn, involves two distinct mechanisms to accumulate sufficient cellular material for regeneration per se — production of new cell mass through cell proliferation and rearrangement of existing cells through cell migration. The relative importance of these two mechanisms remains unclear. Extensive cell division has been documented in virtually all regenerative events in echinoderms studied so far and has been investigated in great detail using thymidine analogs, such as tritiated thymidine, 5-bromo-2’-deoxyuridine (BrdU), and 5-ethynyl-2’-deoxyuridine (EdU) (Leibson, 1992; Candia Carnevali et al., 1998; García-Arrarás et al., 1998; Mashanov et al., 2013; Czarkwiani et al., 2016). On the other hand, the contribution of cell migration to regeneration, has often been merely inferred on the basis of morphological analysis of fixed tissue samples taken at different time points after injury or autotomy (Leibson, 1992; Candia Carnevali, 2006; Mashanov et al., 2005; García-Arrarás et al., 2011; Czarkwiani et al., 2016), but never demonstrated directly. We, therefore, do not know how crucial cell migration is for the success of the regenerative response in echinoderms. Can regeneration be still sustained if cell proliferation is abolished or significantly reduced?

One of the most interesting regenerative phenomena in echinoderms is post-traumatic neurogenesis in their injured CNS. In sea cucumbers, for example, the radial nerve cords (RNCs) fully reconnect and restore their normal cellular architecture within about 3 weeks after complete transection (Mashanov et al., 2008, 2013). Each of the two RNC stumps reorganizes its lesioned surface to form a tubular outgrowth that invades the connective tissue matrix deposited in the wound gap. The two rudiments thus grow towards each other from the opposite sides of the wound and eventually fuse to restore the anatomical connectivity of the injured RNC. One of the most prominent cellular events that unfolds in the lesioned nervous tissue is extensive proliferation of radial glial cells, which has been considered the only major driving force in echinoderm neural regeneration (Mashanov et al., 2013). However, more recent studies of cell turnover in the uninjured adult CNS suggested that some of the newborn cells undergo migration within the neuroepithelium to move away from the place of their birth and populate new locations within the nervous tissue (Mashanov et al., 2015a). We therefore asked if cell migration can also contribute to post-traumatic neural regeneration in echinoderms.

Here, we uncoupled the effect of proliferation from cell migration by using a pharmacological agent, aphidicolin, which inhibits DNA-polymerases of B-family and thus blocks DNA replication in the S-phase of the cell cycle. Unlike many other drugs, which often affect multiple pathways, the effect of aphidicolin is highly specific. In particular, it does not affect cell migration, RNA or protein synthesis. Aphidicolin treatment causes actively cycling cells to pause at the G1/S transition or stop DNA synthesis, if they have already entered the S-phase (Baranovskiy et al., 2014). We demonstrated that, in spite of significant reduction in cell division, the aphidicolin-treated animals successfully formed the outgrowths at the wound surface of their RNC. Surprisingly, those regenerates were of the same absolute size and contained the same number of cells as in the control animals. We concluded that the block of cell division was compensated for by cell migration toward the growing rudiments. We then used vital cell labeling with DiI to show that cell migration was indeed taking place in both the uninjured CNS and in post-traumatic RNC regeneration.

## 2 Materials and Methods

### 2.1 Animal collection and maintenance

Adult individuals of the Caribbean brown rock sea cucumber *Holothuria glaberrima* Selenka, 1867 (Echinodermata: Holothuroidea) were collected by hand from the shallow waters of the rocky intertidal zone of northeastern Puerto Rico (the Old San Juan area). For the duration of the experiment, the animals were kept at room temperature in indoor tanks with aerated natural sea water, which was changed weekly.

### 2.2 Inhibition of cell division in neural regeneration

Aphidicolin was purchased from Sigma Aldrich (A0781) and dissolved in dimethyl sulfoxide (DMSO) to a concentration of 10 mg/mL (0.03 M). This stock solution was stored at −20°C until needed, but no longer than a month. The RNC injury was performed as described elsewhere (Mashanov et al., 2013, 2014, 2015b). Briefly, the animals were anesthetized in 0.2% chlorobutanol (Sigma 112054). The inner side of the body wall was exposed through the cloaca by pushing a glass rod against the epidermis of the ”ventral” mid-body region. The RNC was cut from the coelomic side of the body wall using a sharp razor blade and the animals were returned to the aquaria to regenerate.

To inhibit cell division, we injected aphidicolin at a dosage of 8.3 *μ*g/g (25 *μ*M) into the main body cavity. Each animal received a total of 15 injections. The first injection was done 24 hours post-injury, followed by 14 more daily injections until day 15. The aphidicolin treatment, therefore, covered those phases of regeneration, which involved the formation and growth of the rudiment and highest levels of cell proliferation in the regenerating tissues (Mashanov et al., 2013). These injection parameters were established after a series of pilot experiments, in which a saturation point was reached suggesting that any further increase in drug concentration or in the number of injections will not result in stronger inhibition of cell proliferation. The first injection delivered aphidicolin alone, all subsequent injections also included BrdU (50 mg/kg) to monitor the effect of aphidicolin on the S-phase DNA synthesis. Control animals received DMSO (vehicle) injections. Four animals were injected with the inhibitor and four control animals were injected with the vehicle.

On day 16 of regeneration (i.e., 24 hours after the last aphidicolin/BrdU injection), the tissue samples were fixed overnight at 4°C in 4% paraformaldehyde in 0.01M PBS (pH 7.4), cryoprotected in a series of graded sucrose solutions, embedded in the OCT medium (Sakura), and cryosectioned at 10 *μ*m. Prior to BrdU immunocytochemistry, the cryosections were postfixed for 15 min in formalin vapors. The slides were then rinsed in PBS, pretreated with 0.5% Triton X-100 and soaked in 2N HCl for 30 min at 37°C. After neutralization in 0.1 M borate buffer, autofluorescence was quenched by incubation in 0.1 M glycine at room temperature for 1 hour. The sections were then blocked in 2% goat serum for 1 hour and incubated in a mixture of a primary rat monoclonal anti-BrdU antibody (1:400, GenWay) and the echinoderm glial marker ERG1 (1:1) (Mashanov et al., 2010) overnight at 4°C. After extensive washes in PBS (10 × 10 min), the secondary antibodies (AMCA-conjugated goat anti-rat, 1:50, Jackson ImmunoResearch Laboratories, Inc and FITC-conjugated goat antimouse, 1:100, BioSource) were applied for 1 hour at room temperature. Following the second round of washes (4 × 10 min), the nuclei were stained with propidium iodide and the sections were mounted in an anti-fading medium (2.5% DABCO, 10% Mowiol 4–88 in 25% glycerol buffered with 0.2M Tris-HCL, pH 8.5).

Immunostained sections were photographed with a Nikon Eclipse Ni microscope equipped with a Ds-Qi2 camera using a 40 × objective. Cells were counted in acquired images of every third serial section, at least seven sections per animal, using the CellCounter plugin in the ImageJ/Fiji image analysis software (Schindelin et al., 2012). The cells were counted in the regenerate per se (including the outgrowth and the zone of glial dedifferentiation) plus 100 **μ*m* of adjacent stump tissue, i.e., the region whose general histological architecture did not change after the injury (Fig. 1E, F). The size of the sampling region was measured using standard ImageJ tools.

**Figure 1:**
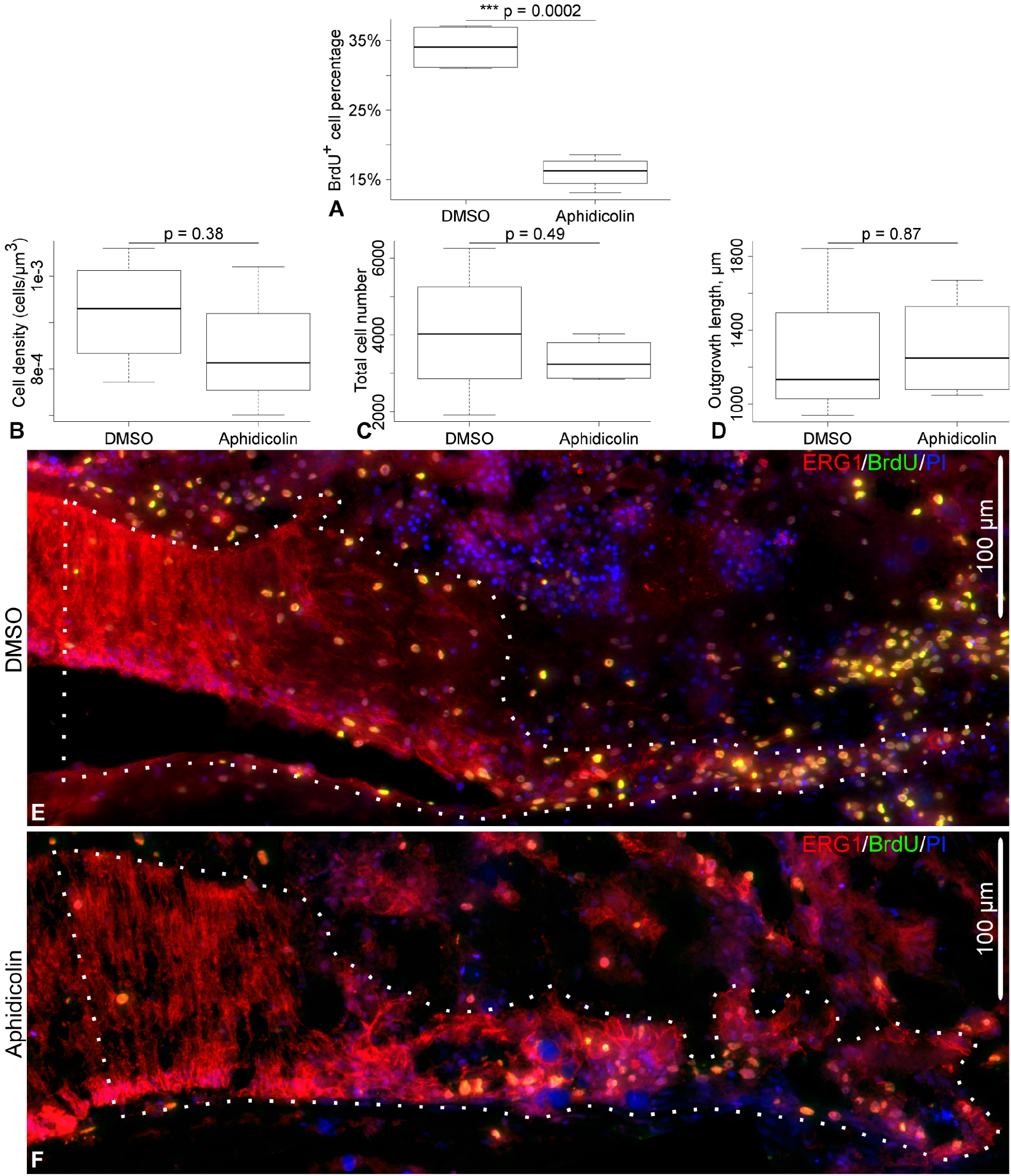
Effect of aphidicolin treatment on radial nerve cord regeneration, day 16 postinjury. (A) The aphidicolin treatment significantly reduced (by a factor or two) the number of BrdU+-cells in the regenerating RNC, but did not affect the cell density (B) in the rudiment, absolute cell number in the rudiment (C), nor the length of the regenerate. (E) and (F) show representative micrographs of the regenerating RNC in a control animal and in an animal injected with aphidicolin, respectively. The growing tip is on the right.

Statistical analysis of the numerical data was performed in R (version 3.3.1) (R Core Team, 2015). The raw data are available in the OpenDocument Spreadsheet (.ods) file format (Additional File 1). The sample R code that can be used to reproduce our statistical computations, as well as the R output are available in Additional File 2. The two-sample Welch t-test was used to test the statistical significance of the aphidicolin treatment on a number of parameters in the regenerating RNC, including the abundance of BrdU+-cells, cell density, absolute cell number, and the length of the outgrowth. The data are represented as whisker plots in Fig. 1A–D, where the box shows the interquartile range (the values between the 25% and 75% percentiles), the thick line in the box is the median, the whiskers are drawn to the data points located within 1.5 × interquartile range outside of the box.

### 2.3 Tracing cell migration with DiI labeling

DiI (1,1’-Dioctadecyl-3,3,3’,3’-Tetramethylindocarbocyanine Perchlorate, ThermoFisher Scientific D282) was dissolved to a final concentration of 2 mg/mL in absolute ethanol. Fine glass needles were hand-pulled from 1 mm diameter glass capillaries. They were dipped into the DiI ethanol solution, the ethanol was allowed to evaporate thus leaving the outer surface of the glass needle covered with a thin layer of DiI. This dye coating procedure was repeated twice.

Two different labeling strategies were used. The first one was designed to label cells at the wound margin. The RNC was injured by complete transection as described above. Immediately after making the incision, the exposed tissues on one side of the wound gap, including the radial nerve cord, were labeled by touching them with a DiI-covered glass needle. The opposite side of the wound was left unlabeled (Fig. 2A, A’). Five animals were sacrificed at each of the two time points after the surgery – on day 2 and day 25 post-injury. The tissue samples were fixed in a mixture of 2% paraformaldehyde and 0.5% glutaraldehyde in 0.01 M PBS (pH 7.4), cryoprotected as above, and cryosectioned at 10 or 20 *μ*m. Sections were collected on gelatinized slides, dried for a few minutes at room temperature and then viewed and photographed without coverslipping (von Bartheld et al., 1990). All sections shown in this paper are longitudinal (i.e., cut along the long axis of the radial nerve cord).

**Figure 2:**
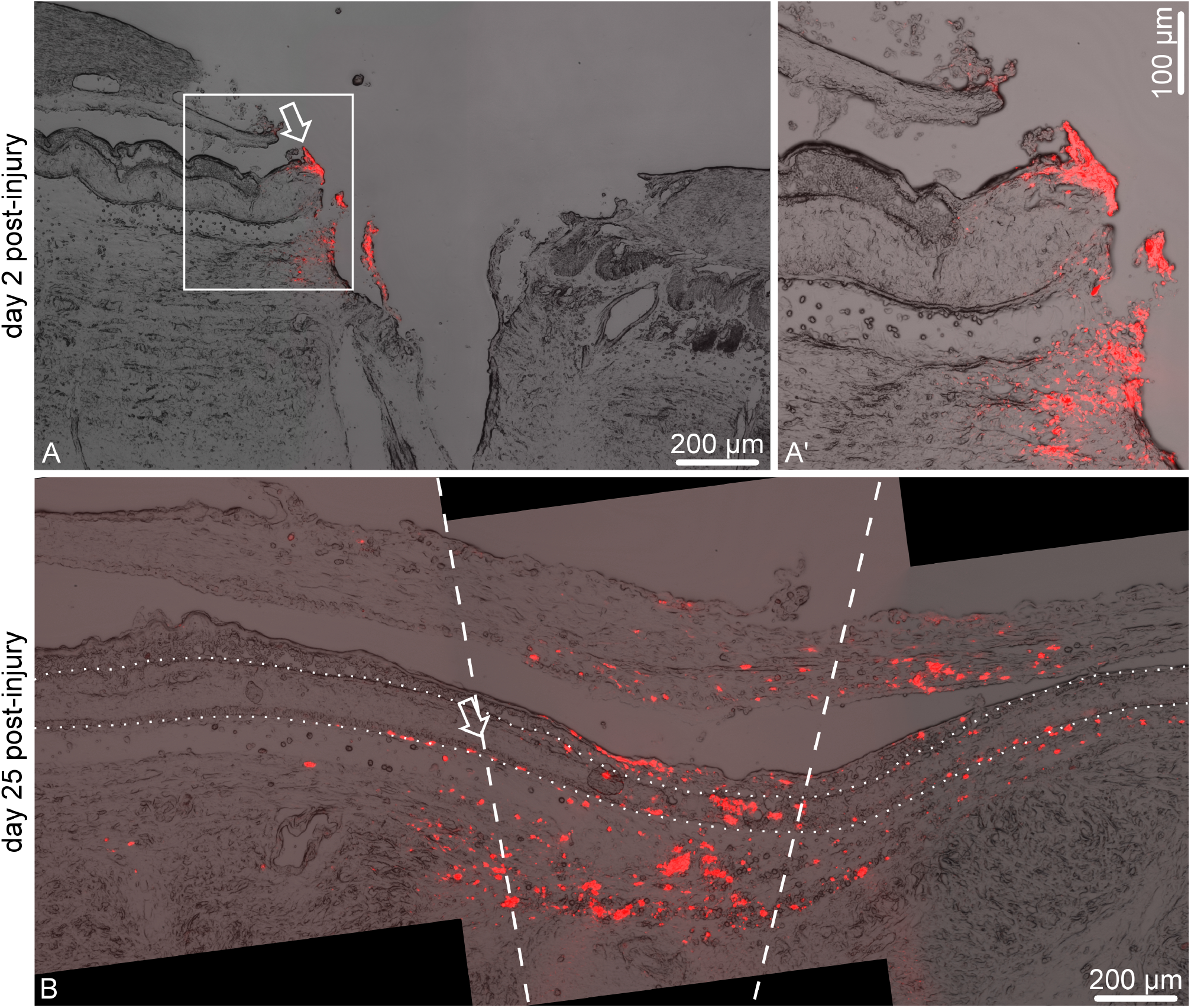
Unilateral DiI labeling of the injured radial nerve cord; longitudinal sections. Immediately after bisection, the dye was applied to one side of the wound, while the opposite side was left unlabeled. The *arrow* shows the site of the original dye application. (A) On day 2 after the surgery, the dye remained on one side of the wound. (A’) Higher magnification of the boxed area in (A). (B) By day 25 after surgery, the wound gap is bridged and the two stumps of the radial nerve cord have reconnected across the wound gap. The *dashed lines* show the original wound margins. Note extensive migration of labeled cells into the new tissue bridging the wound gap. The *dotted line* marks the outline of the radial nerve cord.

The second cell migration tracking strategy involved labeling the cells of the RNC at a distance of about 2 mm away from the wound margin to test if these deeper cells would migrate towards the wound and contribute to regeneration. The animals were anesthetized as above. The radial nerve cord was pricked by a glass needle soaked in DiI solution. The needle was inserted from the inner (coelomic) side of the body wall and, therefore, had to pass trough the coelomic epithelium, radial water-vascular canal, and the radial hemal lacuna before reaching the radial nerve. A single transverse cut was made 2 mm away from the labeling site. The animals were sacrificed on days 2, 16, and 25 after labeling and surgery. At least three animals were used at each time point. The tissue samples were processed, sectioned and analyzed as above. We also included three ”sham” individuals into the experimental design. The RNC of these animals was labeled by piercing with a DiI-soaked needle as above, but was not subjected to transection. These animals were analyzed on day 25 after labeling.

## 3 Results

### 3.1 Aphidicolin reduces cell proliferation in neural regeneration, but does not affect the size of the regenerate

Our previous research indicated a significant increase in cell proliferation that accompanied the growth phase of neural regeneration in sea cucumbers (Mashanov et al., 2008, 2013). It remained unclear, however, whether or not the burst in cell division was the only cellular mechanism involved in formation of the outgrowth at the wound surface of the injured RNC. In order to suppress cell division, we used aphidicolin, an inhibitor of the S-phase DNA synthesis. The treatment regiment was designed so as to continuously inhibit cell division from the early post-injury phase thru the late outgrowth stage. We have previously showed that cell proliferation in the regenerating RNC of H. *glaberrima* starts to increase on days 6–8 post-injury and reaches its peak on days 12–14 post-injury, before returning to the background levels at later stages (Mashanov et al., 2013). For 15 days after injury, the animals were daily injected with either aphidicolin dissolved in DMSO (the treatment group) or with a corresponding amount of DMSO alone (the control group). In addition, to monitor the effect of aphidicolin on the replicative DNA synthesis and thus on cell proliferation, all animals were co-injected with BrdU. As could be expected, aphidicolin treatment resulted in a highly significant 2.1-fold decrease in the abundance of BrdU-positive cells in the regenerating RNC (16.05±1.14% vs 34.06±1.66%, mean±standard error, in aphidicolin-treated and control DMSO-treated animals, respectively) (Fig. 1A, E, F).

Surprisingly, in spite of the significant inhibition of cell division, the regenerating RNC in the aphidicolin-treated individuals had the same total length, absolute cell number and cell density as in the control individuals (Fig. 1B–F). Besides, the growing tips of the regenerating RNCs in both the treatment and control groups showed the same overall tissue architecture (compare Fig. 1E and F) and were composed, as previously described (Mashanov et al., 2008, 2013), of a flattened epithelium made up of dedifferentiated radial glial cells.

How does the regenerating tip of the RNC maintain its size, geometry and cell number when cell division is inhibited? We hypothesized that in addition to cell proliferation *in situ*, there is an additional cell source contributing to regeneration – migrating cells from the more distant regions of the nervous tissue.

### 3.2 Cell migration contributes to radial nerve cord regeneration and also happens in the uninured CNS

In order to study cell migration *in vivo* in the regenerating RNC, we employed vital staining with the lipophilic dye DiI. Two labeling experiments were designed – one to trace the cells at the wound margin (Fig. 2), and the other to label deeper cells located at a distance of about 2 mm away from the plane of injury (Fig. 3A–C).

**Figure 3:**
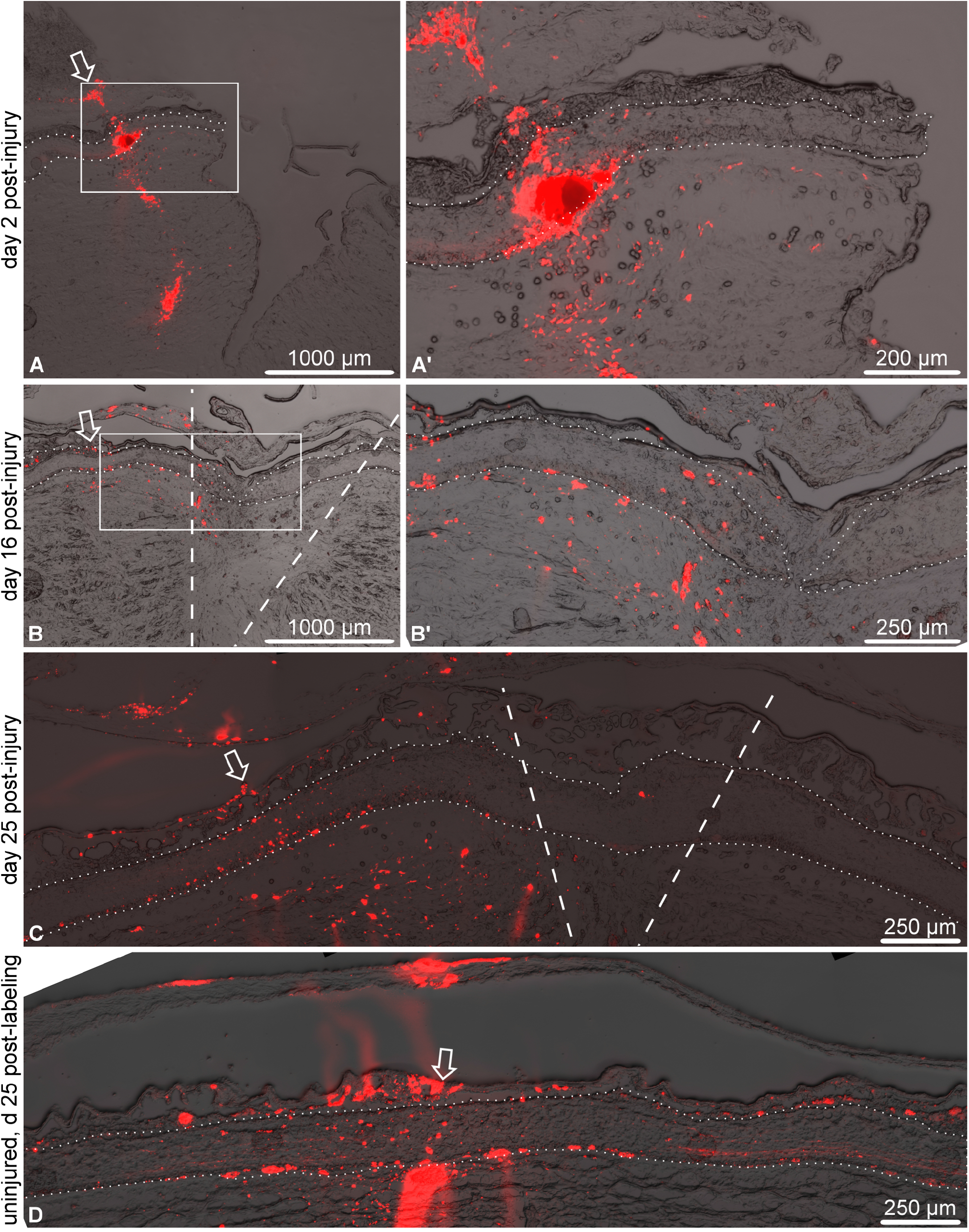
DiI labeling at a distance of 2 mm from the cut on days 2 (A, A’), 16 (B, B’), and 25 (C) after labeling and injury. The site of dye application is indicated by an *arrow.* The radial nerve cord is outlined by *dotted lines. Dashed lines* show the original wound magins. (A’) and (B’) are higher magnification views of the boxed regions in A and B, respectively. (D) Uninjured radial nerve cord 25 days after labeling.

Two days after the surgery and dye application to the cells at the wound margin, five animals were sacrificed and examined to assess the technical success of the first labeling strategy. In all five individuals, DiI-marked cells were seen at one side of the wound only, while the opposite side remained clear (Fig. 2A, A’). By day 25 after injury, the regenerating RNC completely re-connected across the injury gap, and the labeled cells of the wound margin not only have migrated to populate the newly created segment of the RNC, but also moved all the way across the wound gap into the RNC stump at the opposite side of the wound, up to ~1,300 *μ*m from the side of original labeling (Fig. 2B), as measured in cryosections.

The second labeling strategy involved delivery of the dye to deeper cells of the RNC stump, located about 2 mm away from the wound margin (Fig. 3A, A’). Note however that due to fixation and tissue processing, the absolute distances are shorter when measured in cryosections. On day 16 after injury and dye application, the DiI-labeled cells are seen throughout the growing tip of the regenerating RNC (Fig. 3B, B’). In the fully regenerated RNC on day 25 post-injury, many DiI-labeled cells are scattered between the site of dye application and the injury plane and also in the newly created segment of the RNC spanning the injury gap (Fig. 3C), suggesting that deep cells are capable of migrating towards the wound and contributing to RNC regeneration.

We have previously suggested that some cell migration takes place even in the uninjured sea cucumber CNS under normal physiological conditions. That conclusion was based on observations of changes in distribution of BrdU-positive cells in animals left for long chase periods (Mashanov et al., 2015a). Here, we confirmed those earlier observations with vital dye labeling. To this end, we pricked the uninjured mid-ventral RNC at the mid-body level with a DiI-soaked glass needle and kept the animals alive for 25 days. After fixation and cryosectioning, the DiI labeling cells were seen scattered over a distance of over 1,000 *μ*m in both directions along the anterior-posterior axis from the site of dye application (Fig. 3D), suggesting that some cell migration takes place even in the uninjured RNC.

## 4 Discussion

Our results indicate that neural regeneration in echinoderms is a robust and regulative developmental process. In our experiments, a more than two-fold decrease in cell proliferation did not result in a delayed regeneration or in a smaller or deformed RNC outgrowth. On the contrary, the regenerating RNC in aphidicolin-treated animals could still amass the same number of cells and grow to the same absolute size as in the control animals. The only explanation that we can propose is that the deficiency in cell division in the rudiment was compensated for by recruitment of cells through migration.

To directly demonstrate that cell migration in fact takes place, we used the lipophilic vital dye DiI. DiI is irreversibly taken up by cell membranes, without any toxic effect, and, importantly, the labeling does not diffuse from one cell to another (von Bartheld et al., 1990). Two distinct types of cell migration have been implicated in echinoderms tissue regeneration: epithelial invasion, or collective migration of epithelial cells joined by intercellular junctions (Mashanov et al., 2005, 2008, 2013), and migration of individual mesenchymal cells (García-Arrarás et al., 2011; Czarkwiani et al., 2016). So far, it has been proposed that the echinoderm RNC regenerates by reorganizing the cells at the wound site to form a thin tubular outgrowth that later develops into a new RNC segment. The outgrowth is composed of glial cells, which greatly simplify their organization and undergo extensive cell division, but nevertheless maintain intercellular junctions. Epithelial invasion has therefore been proposed as the primary mode of RNC extension across the wound gap. The present study suggest that it might not be the only mechanism. The fact that some of the cells that take up DiI at the wound surface immediately after RNC transection (the first labeling approach, see Results) are found at later stages of regeneration not only in the newly created segment of the RNC that bridges the injury gap, but also within the ”old” former stump RNC region on the opposite side of the wound, suggest that some of the wound surface cells are capable of extensive migration as individual cells. Likewise migration of ”deep” RNC cells from a distance of 2 mm (which equals about 400 neuronal or glial cell body diameters) from the plane of the transection (the second labeling strategy, see Results) towards and beyond the plane of injury can only be accomplished by individual cells that are not anchored to their neighbors by intercellular junctions. Migration of individual cells through the neuroepithelium to new locations, therefore, contributes to redistiribution of cells within the regenerating CNS.

The present study did not clearly establish the identity of the migrating cells. Nevertheless, the available evidence allows us to rule out some of the potential candidates. First, the migrating cells are not mature neurons. For reasons that are still unknown to us, our DiI labeling procedure does not usually yield ”typical” outcome of whole-cell neuronal labeling seen in vertebrates. Only on very rare occasions do we see labeled axonal tracts. Second, due to technical limitations of our surgical procedure we could not avoid labeling some of the mesenchymal cells in the connective tissue regions surrounding the RNC. Even though these connective tissue cells are capable of extensive migration, our previous studies showed that they make no significant contribution to RNC regeneration (Mashanov et al., 2008). The exact nature of the migratory cells in post-traumatic neurogenesis in echinoderms remains an important question to be resolved by future studies.

In spite of the chronic exposure to aphidicolin, the cell division in the RNC regenerate, although decreased more than two-fold, did not stop completely. We do not think that the inability of our aphidicolin treatment to completely abolish cell division was due to insufficient concentration of the drug or the duration of the procedure. First, the dosage that we used (25 *μ*M) was similar or higher than the range of concentrations used in similar experiments with echinoderm and vertebrate embryos, larvae, organ and cell cultures (0.6–30 *μ*M), which achieved complete or near-complete cell division inhibition (Fishman et al., 2015;

Geling et al., 2003; Krupke and Burke, 2014; Smith et al., 2009; Shang et al., 2010). Second, our preliminary experiments showed that treatments at lower doses and smaller number of injection caused a similar two-fold decrease in mitotic activity in RNC regeneration. The increase in dosage and in the number of injection, however, resulted in much smaller variation in response to the treatment between individuals. Third, even at much higher concentration (150 *μ*M) aphidicolin did not completely stop cell division in *Xenopus* embryos. The authors suggested that this observation was due to some residual DNA synthesis, which allowed the cells to complete the S-phase and go into the mitotic cycle (Harris and Hartenstein, 1991). Depending on the specific properties of an organism, cell division inhibitors are known to vary in extent of their effect on cell proliferation. For example, one of the factors that prevents the general use of aphidicolin as an anticancer drug in human patients is its very rapid clearance from the human plasma (Baranovskiy et al., 2014). The partial resistance of the regenerating echinoderm nervous tissue to aphidicolin treatment may provide some interesting insights into DNA replication biology and physiology of echinoderms.

The present study, therefore, directly demonstrates that migration of individual cells plays an important role in post-traumatic regeneration in echinoderms and presumably can rescue regeneration response if cell proliferation is suppressed. This report adds to the growing realization that migration of individual cells is a significant, although traditionally underappreciated and often improperly studied, factor contributing to post-embryonic developmental phenomena, such as post-traumatic regeneration and normal cell replenishment, in a wide range of multicellular animals (Zattara et al., 2016).

## 5 Conclusions

- Chronic aphidicolin treatment resulted in a significant decrease in cell proliferation in the regenerating RNC of the sea cucumber *H. glaberrima*
- The RNC in the aphidicolin-treated animals, however, continued to regenerate and did not differ from the vehicle-treated controls in size, internal structure, or cell number.
- We speculate that the inhibition of cell division was compensated for by cell migration from more distant regions of the neuroepithelium located as far as 2 mm from the site of the original injury and outgrowth formation.

## 6 List of Abbreviations

*BrdU* –: 5-bromo-2-deoxyuridine;
*DiI* –: 1,1’-Dioctadecyl-3,3,3’,3’-Tetramethylindocarbocyanine Perchlorate;
*DMSO* –: dimethyl sulfoxide;
*RNC* –: radial nerve cord

## 7 Acknowledgments

The study was supported by grants from the NIH (1R03NS065275-01) and the NSF (IOS-0842870, IOS-1252679), as well as by the University of Puerto Rico

## 9 Additional Files

### Additional File 1

OpenDocument Spreadsheet (.ods) file containing the raw measurements of BrdU+-cell abundance, total cell number and cell density in the radial nerve cord regenerate, as well as the outgrowth length. This file can be opened with/imported into all major spreadsheet tools, including LibreOffice/OpenOffice Calc and MS Office Excel.

### Additional File 2

R code used to perform the statistical computations and generate the box and whisker plots

